# PiggyBac mediated transgenesis and CRISPR/Cas9 knockout in the greater waxmoth, *Galleria mellonella*

**DOI:** 10.1101/2024.09.17.613535

**Authors:** James C. Pearce, Jennie S. Campbell, Joann L. Prior, Richard W. Titball, James G. Wakefield

## Abstract

The larvae of the greater waxmoth, *Galleria mellonella*, are gaining prominence as a versatile non-mammalian *in vivo* model to study host-pathogen interactions. Their ability to be maintained at 37°C, coupled with a broad susceptibility to human pathogens and a distinct melanisation response that serves as a visual indicator for larval health, positions *Galleria* as a powerful resource for infection research. Despite these advantages, the lack of genetic tools, such as those available for zebrafish and fruit flies, has hindered development of the full potential of *Galleria* as a model organism. In this study, we describe a robust methodology for generating transgenic *Galleria* using the PiggyBac transposon system and for precise gene knockouts via CRISPR/Cas9 technology. These advances significantly enhance the utility of *Galleria* in molecular research, opening the way to its widespread use as an inexpensive and ethically compatible animal model for infection biology and beyond.

## Introduction

The larval stage of the greater waxmoth, *Galleria mellonella*, is increasingly recognised as a valuable *in vivo* mammalian replacement model, particularly in the fields of infection, immunology, and inflammation ^1–9^. They undergo melanisation in response to immune challenges and have broad susceptibility to a range of medically important microbes (reviewed in Asai et al 2023,Kavanagh and Sheehan 2018, Menard 2021, Kavanagh & Fallon 2010, giammarino et al 2024, Tsai et al 2016 ^1–3,5,10,11^). Their capacity to be maintained at 37°C confers a significant advantage over other model systems such as fruit flies or zebrafish, particularly for studies involving human pathogens. Moreover, unlike vertebrate models, *Galleria* larvae are not subject to stringent regulatory or licensing requirements. Finally, recent discoveries, such as their unique ability to metabolise polyethylene and polystyrene independent of their microbiota ^12–15^, could yield advances in our understanding of plastic degradation and solutions to plastic waste and underscore the potential for broad application of *Galleria* larvae in research settings.

The availability of multiple *Galleria* genomes, first published in 2018 ^13,16–18^, have resulted in an and expanded set of molecular and cellular tools, alongside transcriptomic and proteomic datasets ^19–28^. These resources have significantly increased the potential of *Galleria* to be developed as an alternative to mammalian infection models. However, the absence of robust genetic manipulation techniques – critical for the insertion, deletion and engineering of genes – remains a limiting factor. While such techniques have been extensively deployed for other insects, the application of transgenic and genetically modified (GM) approaches in *Galleria* have yet to be developed.

Among the most widely adopted tools for genetic modification are the PiggyBac transposase system and CRISPR/Cas9, both of which offer powerful, complementary means of creating transgenic organisms and targeted gene knockouts. The PiggyBac transposase system, isolated from *Trichoplusia ni* ^29^, allows for the seamless integration of genetic material into TTAA nucleotide sequences across a wide array of animal species, facilitated by a separately provided transposase enzyme ^30,31^. The ability of this method to insert large genetic cargos ^32^ without the need for specific landing sites, while advantageous, also poses challenges, including variable integration efficiency ^33^ and the potential for disruption of endogenous gene function if insertion occurs within a coding region. CRISPR/Cas9-mediated mutagenesis, has rapidly become the gold standard for targeted genetic modification, surpassing other techniques such as Zinc Finger nucleases (ZFNs) and transcription activator-like effector nucleases (TALENs) ^34–36^. This technique employs a bipartite type II CRISPR system to direct a CRISPR associated nuclease (Cas) to specific genomic loci via an RNA guide with a complementary base sequence ^37–39^. The resultant double strand breaks (DSBs) are repaired with varying fidelity by different DNA repair pathways, enabling either disruption of endogenous gene function or the insertion of exogenous genetic sequences ^40^.

In this study, we successfully apply both the PiggyBac and CRISPR/Cas9 systems to *Galleria mellonella*, demonstrating their efficacy in generating transgenic lines and gene knockouts. These advances significantly enhance the genetic tractability of *Galleria*, establishing a foundation for its broader application across diverse research domains.

## Materials and Methods

### Animals & Rearing

An inbred “wild type” (WT) *Galleria mellonella* colony was reared on an artificial honey diet (Diet 3, Jorjao et al. 2018^41^) at 30 °C, constant darkness. A more detailed rearing protocol is described in Pearce et al. (2024).

Transgenic strains were reared at 30 °C on the same diet in large polypropylene fly vials (10 cm x 5 cm – Darwin Biological) with foam bungs to prevent L1 larval escape. At the late larval wandering stage, larvae were transferred to petri dishes containing diet and allowed to pupate. Pupae were transferred to small PET jars with 5 μm wire mesh lids and the adults allowed to oviposit on egg papers.

### Histology

Jars of WT adults were kept at 30 °C in constant darkness and allowed to lay on egg papers overnight. The egg papers were removed, and the embryos discarded before replacing the clean egg papers into the jars. The moths were allowed to oviposit undisturbed for 1 hr in darkness before the papers were again removed and the embryos collected and labelled as 0-1 hr old. Embryos were allowed to develop for set time periods before dechorionating for 2 min in a diluted solution of thin bleach (1.25 % active chlorine) and 0.05 % Triton X-100.

Aged embryos were transferred into a 1.5 ml Eppendorf containing 500 μl of heptane and 500 μl of methanol, inverted several times until the majority of the embryos at the interphase of the two solutions dropped into the methanol. The heptane and any embryos remaining at the interphase were removed using a Pasteur pipette, before washing twice in fresh methanol. Embryos were stored at 4° C for no more than a week, until use.

Embryos were rehydrated sequentially for 15 min each in 75:25 and 50:50 methanol:PBS + 0.01 % Triton X-100 (PBST), before further 15 min rehydration in PBST. They were stained with 0.5 mg/ml Hoechst 33258 in PBST for 20 min at room temperature, before final 3 x 5 min washes in PBST.

Larval tissues were fixed in 4% paraformaldehyde/PBS + 0.1% Tween for 1 hour and stained overnight with a rabbit anti-GFP polyclonal antibody (Abcam 6556) and labelled using an appropriate Alexa Fluor 488 secondary dye (Molecular Probes).

### Imaging

Embryos were mounted on microscope slides, within 2 stacked ring binder reinforcement stickers, in Vectashield mounting media (Vectalabs) and imaged using a Nikon TE-2000U inverted microscope. Larvae were anaesthetised using CO_2_ and imaged under a Leica MZ10F fluorescence stereomicroscope with a GXCAM HiChrom-HR4 HiSens camera. Fixed larval tissues were imaged using a Leica SP-8 confocal microscope.

### RNA/DNA Extraction & Polymerase Chain Reactions

*Galleria* tissue samples were flash frozen in liquid nitrogen and homogenised in Eppendorfs using disposable microcentrifuge pestles (DWK life sciences), then stored on ice until use.

Total RNA was extracted from homogenates of 0-6 hr embryos and the posterior 1/3^rd^ larvae (larval end segment) using a Trizol reagent/chloroform extraction according to manufacturers protocol and ethanol precipitated before either being used immediately or stored at -80 °C. cDNA was generated using a High Capacity cDNA reverse transcription kit (Applied Biosytems) using the manufacturer’s protocol.

Genomic DNA was extracted from whole or partial tissue samples, depending on the size, using the New England Biolabs (NEB) gDNA extraction kit with NEB’s insect tissue protocol.

DNA fragments for diagnostic purposes were amplified using a GoTaq Hot Start mastermix (Promega). For all other purposes (including sequencing), fragments were amplified using a KOD Hot Start polymerase kit (Toboyo), according to manufacturer’s instructions.

### Plasmids

Plasmids pHA3PIG ^42^ and pBAChsp90GFP-3xP3DsRed ^43^ were a kind gift from Professor Hideki Sezutsu. Henceforth we will refer to pBAChsp90GFP-3xP3DsRed as p*Bm*hsp90:GFP/3xP3:DsRed to differentiate between the different *Galleria* and *Bombyx* hsp90 promoters. All plasmids were assembled using Gibson assembly.

p*Bm*hsp90:hyPB was generated by inserting the Bombyx mori hsp90 2.9 kb fragment from p*Bm*hsp90:GFP/3xP3:DsRed and the hyper-active Piggybac transposase from SPB-DNA (Hera Biolabs) upstream of wild type PiggyBac transposase 3’ UTR from pHA3PIG in the digested backbone of pHA3PIG.

p*Gm*hsp90:GFP-αtub1b was generated by creation of an expression cassette consisting of the 2 kb upstream region of *Galleria* hsp83 (hsp90) placed directly upstream of an N-terminal eGFP tagged *Galleria* α tubulin 1b cDNA sequence (LOC113521067) and SV40 polyA terminator, which was inserted between the two PiggBac inverted terminal repeats of the digested p*Bm*hsp90:GFP/3xP3:DsRed backbone.

p*Bm*hsp90:histone2av-Mch/3xP3:DsRed was generated by digesting the p*Bm*hsp90:GFP/3xP3:DsRed plasmid with PmlI and AscI and inserting a synthesised fragment consisting of the *Galleria* Histone 2AV sequence (LOC113518755) connected to mCherry, via a short linker with a SV40 poly A terminator.

All plasmids were propagated in NEB 10β cells and midi-prepped using a Nucleobond Xtra midi kit (Macherey-Nagel). They were ethanol precipitated before reconstituting in either nuclease free water or filter sterilised 5 mM phosphate/5 mM KCl injection buffer, pH 7.4.

### Embryo Microinjection

The injection protocol is described in detail in Pearce et al. (2024), but briefly 1-2 hr old embryos (stored at 30 °C) were dechorionated using a diluted bleach solution and aligned along the edge of coverslips, glued to glass slides.

Injection mixes were made by adding plasmids, guides or proteins sequentially to injection buffer and checking total concentration using a nanodrop. Before use, injection mixes were spun at 16,000 g for 1min, before gently aspirating off the upper 90% and transferring to a fresh Eppendorf tube. The spun mixes were then stored on ice until use.

Embryos were injected using an Eppendorf Injectman 4 microinjection system mounted to a Nikon Eclipse TE2000-U inverted microscope with a small volume of injection mix, equal to a droplet roughly 1/5 diameter of the embryo. For Piggybac-mediated mutagenesis, embryos were injected between 3-5 hrs post oviposition (PO) and the injection droplet was placed on the side of the embryo relative to the putative anterior-posterior poles. For CRISPR/Cas9, embryos were injected at 2-3.5 hrs PO and injection droplets placed in the centre of the embryo.

### Post-injection rearing and screening

Post injection, embryos were reared as described in Pearce et al (2024). Late stage G0 larvae were removed from diet and screened for visible changes in somatic fluorescence using a fluorescence stereo microscope (Leica MZ10F or Olympus SZX-16). Those demonstrating mosaic fluorescence were separated into fresh diet. Once pupated, putative transformants were sexed, separated the basis of genital morphology visible in the terminal segments, and G0 adults mated to either a mixture of siblings and WTs (PiggyBac injected G0s) or WTs only (CRISPR/Cas9 injected G0s). Embryos were collected from each cross and the progeny screened for visible changes in somatic fluorescence at both embryonic and larval stages. Stable transgenic lines were generated by selecting the G1 generation with the brightest phenotype and outcrossing for 2-3 further generations, before sibling mating and screening for the brightest offspring. These were then sibling mated and screened for consistent phenotype over multiple generations.

### Guide RNA synthesis and Cas9

An anti-eGFP sgRNA previously described by Jao et al ^44^ was synthesised *in vitro* as described in Burger et al. ^45^. A DNA template for the sgRNA was assembled by PCR amplifying two primers, sgRNA-EGFP fwd and a PAGE purified sgRNA rev, and then the product purified (Promega). *In vitro* transcription was performed overnight at 37 °C using a T6 RNA polymerase (Roche), before DNase treatment, RNA clean-up and validation for size and presence of a single band on a denaturing MOPs gel. The sgRNA was stored at -80 °C until use.

The Cas9 protein used was a kind gift from Professor Christian Mosimann and is a modified version of *S. pyogenes* Cas9 fused in frame with an additional C-terminal HA tag, a bipartite nuclear location sequence (NLS), an mCherry polypeptide sequence and an additional monopartite NLS at the very C-Terminus ^45^.

### Mutation analysis & Sequencing

Piggybac insertion sites were verified via inverse PCR. gDNA from G1 transgenic larvae was digested for 2 hrs using HpaII, before heat inactivation. Genomic fragments were then self-ligated using T4 DNA ligase at 4 °C overnight, before ethanol precipitating. The genomic regions surrounding the Piggybac entry sites were amplified through PCR using two sets of primers specific to the 5’ and 3’ Piggybac ITRs (iPCR5’ F & R and iPCR 3’ F & R), before sequencing using a commercial short read service (Eurofins). Mutations in the GFP coding region were amplified using primers for GFP (GFPF & GFPR) and sequenced using a commercial short read service (Eurofins).

## Results

### Embryonic development timings indicate a 6hr window for microinjection

*Galleria* were reared at 30 °C and embryos were deposited on egg papers, prior to fixing. Imaging of early embryos, stained for DNA, revealed developmental timings postoviposition. In embryos aged between 1.25 – 2.75 hrs, postoviposition, sperm nuclei could be observed in transit towards the ova nucleus, with polar bodies migrating towards the boundary of the embryo (**Figure 1A-C**). A small proportion of embryos in this time window were also observed to have undergone their first mitotic division; with some beginning their second.

**Figure 1:**
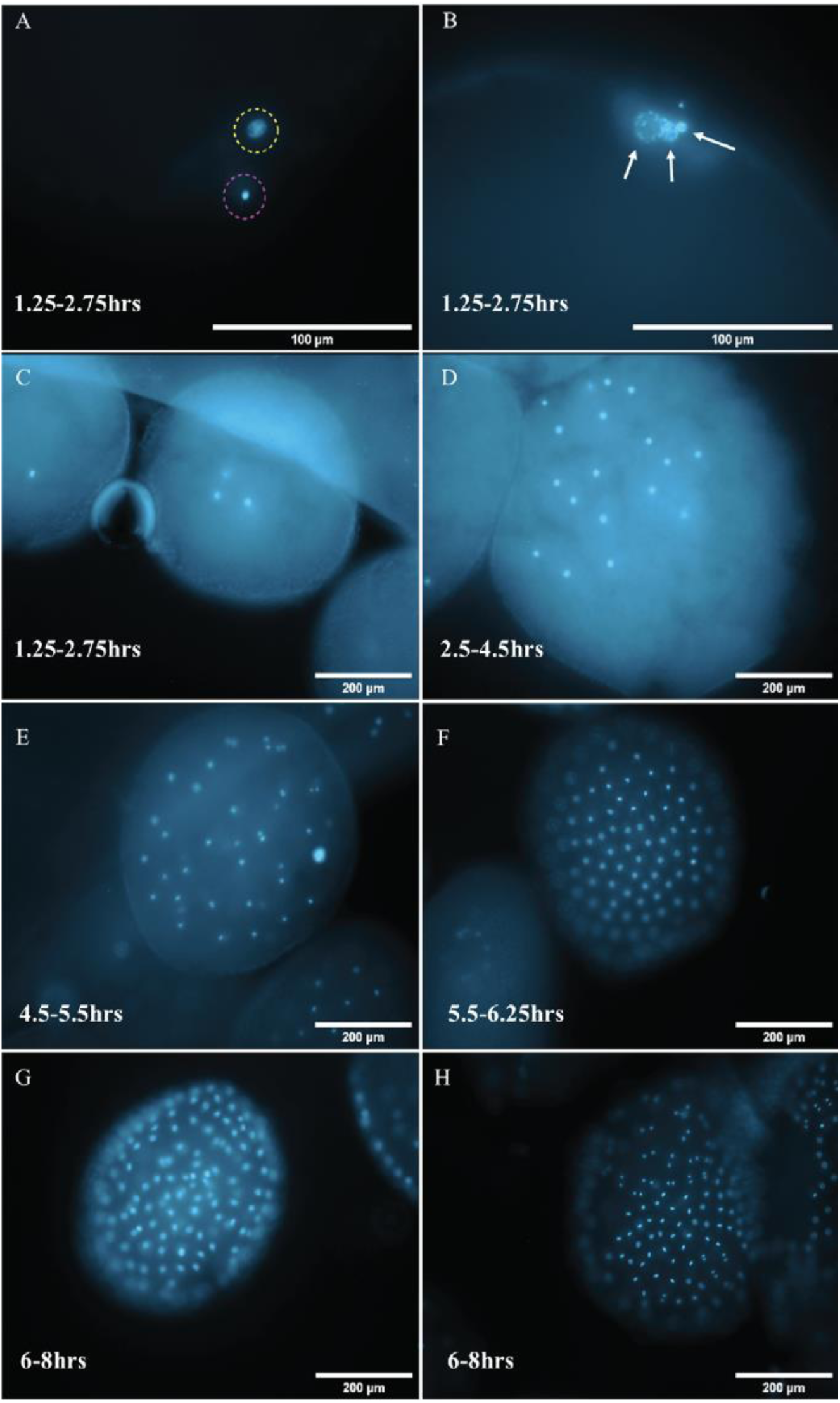
*Galleria* embryos fixed during set points in development and stained with Hoechst 33258 DNA dye. In the first 1.25-2.75hrs (Images A-C) post oviposition (PO) the sperm nuclei (Image A— purple circle) can be seen moving towards the ovum nuclei (yellow circle) and nuclei resembling polar bodies gather towards the periphery of the embryo (Image B—white arrows). This timeframe appears to cover up to the second mitotic division (Image C). From 2.5-5.5hrs PO, energids migrate towards the periphery with a bias to the anterior pole (Image D) and by 4.5-5.5hrs PO the first ones have just reached the periphery (Image E). Nuclei are dividing synchronously at this point as they share a common cytoplasm. All energids have reached the periphery by 6.25hrs and are still dividing together (Image F), although synchronicity begins to be lost from 6-8hrs (Images G & H) potentially indicating the onset of cellularisation.

In batches of embryos collected within the next 3.5 hours (2.75-6.25 hrs post oviposition) increased numbers of nuclei, migrating toward the periphery of the embryo were observed (**Figure 1D-F**). The nuclei in different embryos appeared to be at the same cell cycle stage, as determined by chromosome condensation, alignment and segregation, indicating that, at this time point, they still share a common cytoplasm (**Figure 1D-F**). This synchronicity was lost however, in batches of embryos fixed and imaged 8 hrs post oviposition, indicating that cellularisation occurs between 6.25-8 hrs (**Figure 1G & H**). As embryos developed further, a difference in nuclei spacing became noticeable, with the future embryonic tissue being more densely nucleated by 14 hrs (not shown). Together, this analysis provides a time window for injection to generate stable germline transformants of between 0-6 hrs of development at 30 °C.

### p*Bm*hsp90:hyPBase but not pHA3Pig is a suitable as a donor plasmid for PiggyBac mutagenesis in *Galleria*

To unequivocally inject exogenous material prior to cellularisation, we performed manipulations on 0-2 hr old dechorionated embryos (see Methods). Perhaps unsurprisingly, injection with piggyBac DNA plasmids caused mortality (**Table 1**), with between 6-19 % of injected embryos surviving to pupation. Where performed, hatch rates were much lower than injections with only injection buffer (Pearce et al., 2024). However, given the large number of embryos that can be collected and injected within a few hours we proceeded with this methodology to screen for transformants.

**Table 1:**
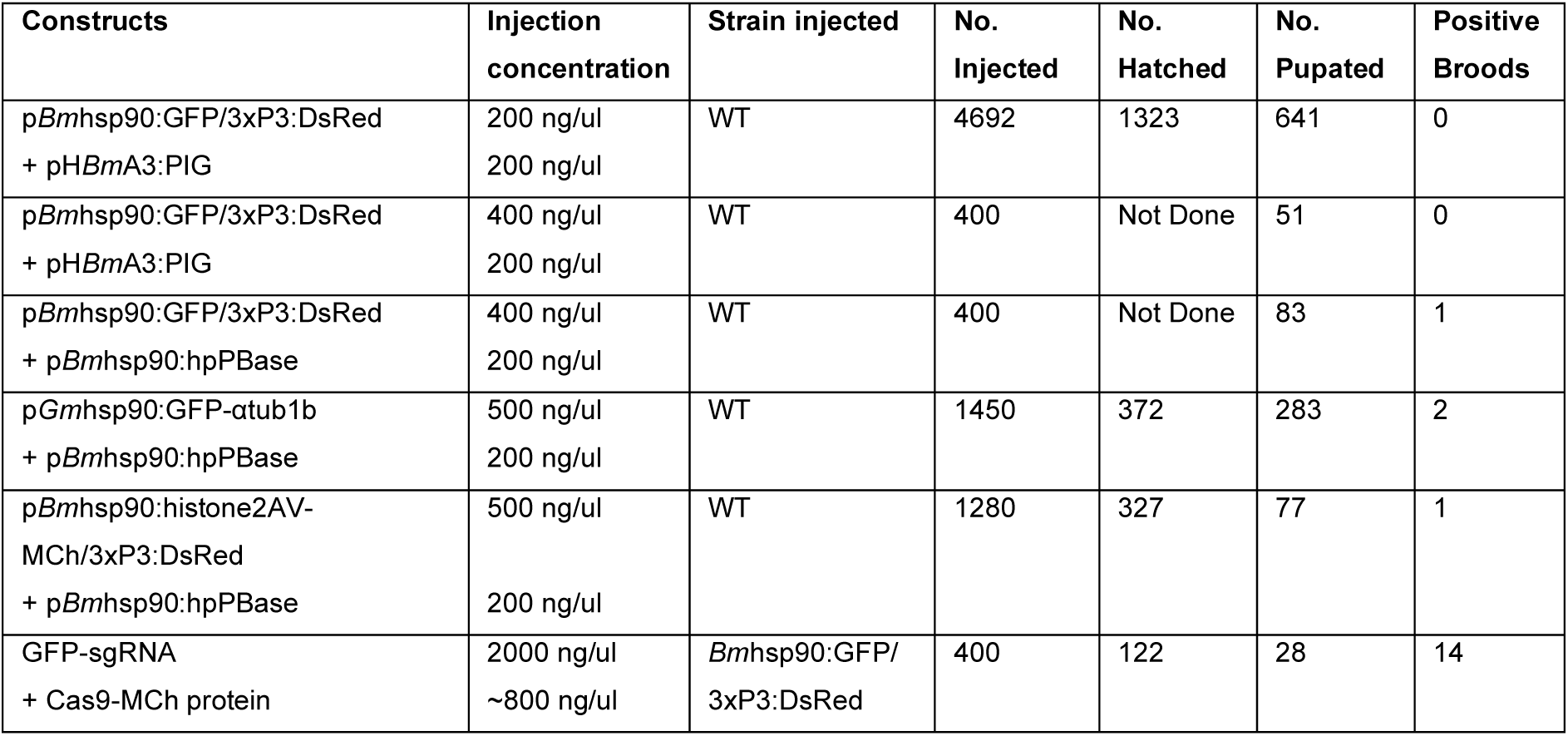
Hatch and pupation rates for different injection mixes. Where hatch rates were not calculated, Not Done has been entered.

Initially, we injected a 200:200 ng/μl plasmid mix consisting of pBAChsp90GFP:P3dsRed donor and pHA3Pig helper, as used by Tsubota et al ^43^. This plasmid was expected to drive GFP ubiquitously (under the control of the hsp90 promoter) and dsRed in the developing eye (under the control of the P3 promoter). However, we did not observe any GFP or dsRed fluorescence in any G0 or G1 *Galleria*, despite a large number of embryonic injections (N=4692) and G1 broods screened (N>600) (**Table 1**). This indicates either that this helper plasmid has very low activity in *Galleria*, or that the promoters are inefficiently driving fluorescent protein expression.

To address these issues, two sets of injections were performed with the same donor, p*Bm*hsp90:GFP/3xP3:DsRed, but at a higher concentration (400 ng/μl), either with 200 ng/μl of the original pHA3:PIG helper or with a different helper plasmid, p*Bm*hsp90:hyPBase, which encodes a hyperactive PiggyBac transposase mutant, driven by the same Bombyx hsp90 upstream sequence as used in the donor ^44^ (**Figure 2A**). Mosaic green fluorescence was observed in several G0 larvae from the p*Bm*hsp90:hyPBase injection group, while none was observed in the pHA3:PIG group (**Table 1**). From the p*Bm*hsp90:hyPBase group, eGFP and DsRed positive transgenic larvae were recovered from the brood of a single G0 adult – WT cross (**Table 1**).

**Figure 2:**
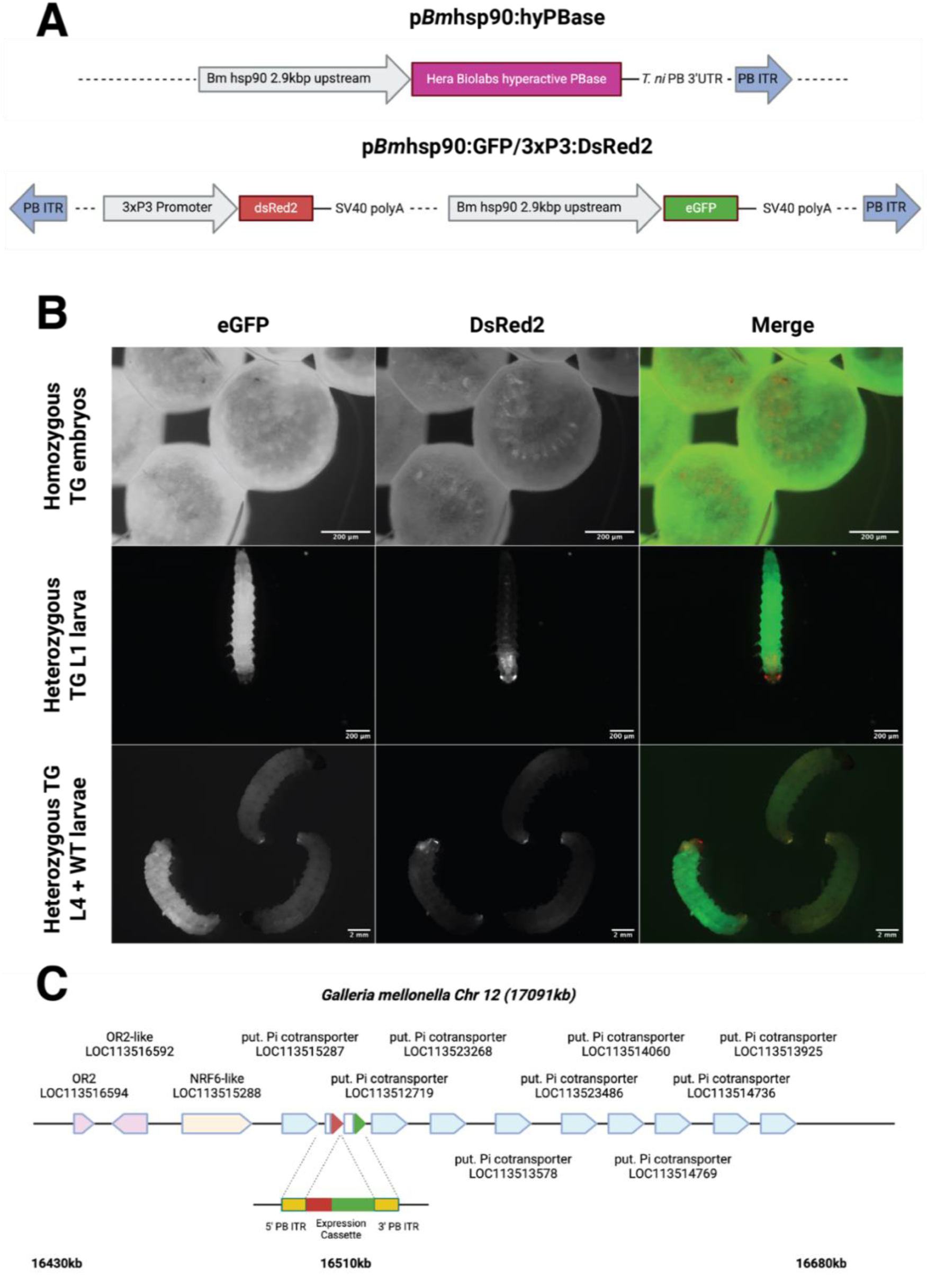
The helper and donor constructs used to develop *Bm*hsp90:GFP/3xP3:DsRed transgenic strain (A). Both eGFP and DsRed expression can be observed in the late embryo (B, top row), first instar larvae (B, middle row) and 4^th^ instar larvae (B, bottom row). eGFP expression is very bright within the unmelanised tissues, with DsRed expression limited to neural tissue including the brain, optic nerves and stemmata. The transgenic cassette appears to have within an inorganic phosphate co-transporter gene cluster on chromosome 12, adjacent to a small cluster of odorant-like receptors.

### The *Bombyx mori* hsp90 promoter sequence drives strong but not constitutive activity in Galleria and 3xP3 promoter appears to be neural specific

In larvae transformed with p*Bm*hsp90:GFP/3xP3:DsRed, bright eGFP expression could be observed in larval, pupal and adult stages. However, instead of ubiquitous expression, eGFP in embryos appeared to be limited to the vitellophages and was absent both from the germ band and developing nascent larva until just before eclosion (**Figure 2B**). In larvae, eGFP expression was strongest in the muscle, fat body and malpighian tubules with weaker expression seen in the gut, silk glands and epidermis.

As expected, DsRed expression was observed to be present in, and solely within, neural tissue, with strongest expression in the eyes, optic nerves, brain and segmental ganglia (**Figure 2B**). It was first observed at around day 5 of embryonic development at 30 °C, at the anterior and in a segmented pattern within the embryo, perhaps corresponding to the developing eyes and segmental neural centres.

To map the insertion site in one of these transformants, inverse PCR was undertaken. The flanking regions of the *Bm*hsp90:GFP/3xP3:DsRed expression cassette were found to correspond to the proximal end of chromosome 12, within an intergenic region between LOC1131515287 and LOC1131512719, in a cluster of putative inorganic phosphate cotransporters (Fig 2 C). Visual monitoring of outward health and developmental timings for this transgenic line, across multiple generations, showed no difference to WT, suggesting that transgene expression at this locus is not deleterious to organism.

### Generation of *Galleria* α tubulin and histone cellular reporter lines

To investigate whether a *Galleria* hsp90 promoter might drive constitutive expression, and whether PiggyBac would be suitable for creating reporters of cellular and subcellular dynamics, two DNA constructs were generated. In the first (*Gm*hsp90:GFP-αtub1b), the *Galleria* α tubulin 1b gene, one of several α tubulin genes in the *Galleria* genome, was N terminally tagged with eGFP and placed downstream of a 2 kb region corresponding to the *Galleria* hsp90 promoter (**Figure 3A**). In the second, a 2kb region corresponding to the Bombyx mori hsp90 promoter (Kawasaki et al 2003) was placed upstream of the *Galleria* monocistronic histone 2A variant, with a C terminus mCherry tag (*Bm*hsp90:his2av-MCh) (**Figure 3A**).

**Figure 3:**
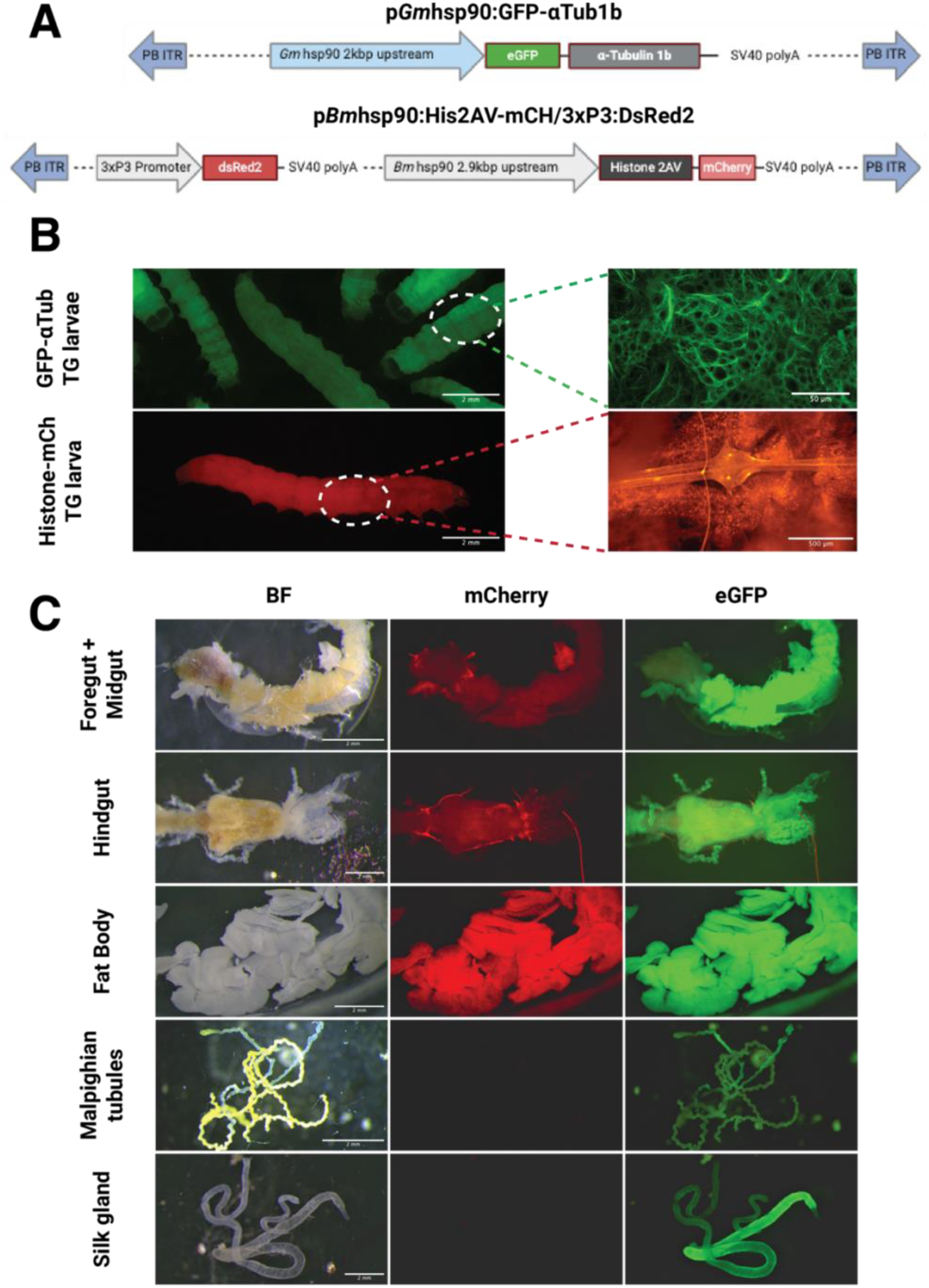
Constructs used to generate transgenic lines *Gm*hsp90:GFP-αtub1b and *Bm*hsp90:his2av-MCh (A). *Gm*hsp90:GFP-αtub1b (B, top left) with fat body tissue fixed and stained with an anti-GFP antibody (B, top right) and *Bm*hsp90:his2av-MCh (B, bottom left) larvae with fat body and dorsal neural ganglion imaged live (B, bottom left). Expected cystoskeletal distribution of eGFP was observed, corresponding with expected localisation of tubulin, with a nuclear localisation observed for mCherry. Brightfield (BF), mCherry and eGFP tissue expression patterns in a strain with both *Bm*hsp90:his2av-MCh/*Gm*hsp90:GFP-αtub1b expression cassettes (C). Strong fat body expression was observed for both fluorophores, with the *Galleria* hsp90 promoter appears to drive strongest in gut and silk gland and the *Bombyx* hsp90 promoter stronger in epidermal and muscle tissue (not shown) but very weak in silk glands and malpighian tubules.

Hatch rates for injected embryos were similar for both constructs, at around 26% survival (**Table 1**). However, while 20% of *Gm*hsp90:GFP-αtub1b injected embryos reached pupation, this proportion was 6% for embryos injected with *Bm*hsp90:his2av-MCh (**Table 1**). This may indicate some additional toxicity due to integration of the histone expression cassette, either due to overexpression of this histone variant or the insertion locus itself.

Transgenic lines for both constructs were obtained. Although inverse PCR failed to locate the insertion locus for either line, fluorescence corresponding to microtubules and nuclei, respectively, could be observed in embryos (not shown) and larvae (**Figure 3B**). Interestingly, we found differences between the expression pattern of the *Bombyx* and *Galleria* hsp90 integrated lines. Strong eGFP expression for the *Galleria* hsp90-GFP-αtub1b construct was observed in the fat body, hindgut, midgut and silk glands, with somewhat weaker expression in the malpighian tubules (**Figure 3C**), as well as the muscle and epidermis (not shown). In contrast, the *Bombyx* hsp90 promoter drove expression of mCherry-his2av strongly in the fat body, more weakly in the gut, but appeared absent, or very low levels, in the silk glands and malpighian tubues (**Figure 3C**). This is in contrast with results previously observed for eGFP expression in the *Bm*hsp90:GFP/3xP3:DsRed strain, where very strong expression was observed in malpighian tubules and moderately in silk gland; suggesting either insertion-specific differences or selective suppression of His2av expression in those tissues. Embryonic expression for both promoters was weak, although some germ activity could be observed for the *Galleria* hsp90 vs the vitellophages in the Bombyx hsp90 (not shown).

### CRISPR/Cas9 mediated mutagenesis in Galleria

Although PiggyBac transgenesis is a useful tool, we sought to further extend the molecular engineering capabilities of *Galleria* through testing the efficacy of CRISPR/Cas9, with respect to gene knockouts. An injection mix consisting of a KCl-buffered ribonucleoprotein complex of single sgRNA (in molar excess) targeting the eGFP sequence ^44^ and mCherry tagged Cas9 was injected into embryos, homozygous for the *Bm*hsp90:GFP/3xP3:DsRed transgenic cassette. A hatch rate of 31% was similar to that observed for PiggyBac injections, yet survival to pupation was low at only 7% (**Table 1**).

Within the developing G0 larval offspring, a range of eGFP knock-out phenotypes were observed, varying from minor fluorescence mosaicism to an almost complete lack of fluorescence in somatic tissue (not shown). All G0 adults were outcrossed to WT mates, and the resulting broods screened for eGFP-negative G1 larvae that were positive for DsRed expression in their stemmata (**Figure 4A**). 50 % (14/28) of broods contained such knock-out larvae, including offspring from G0 parents that had shown no or very minor loss of GFP expression. Analysis of the resulting CRISPR mutants via Sanger sequencing revealed a combination of small indels and larger deletions around the guide target site (**Figure 4B**). Off-target effects were not screened for; instead crispant larvae were outcrossed to WT strains for 3 generations before creating a stable line, thus minimising accumulated mutations.

**Figure 4:**
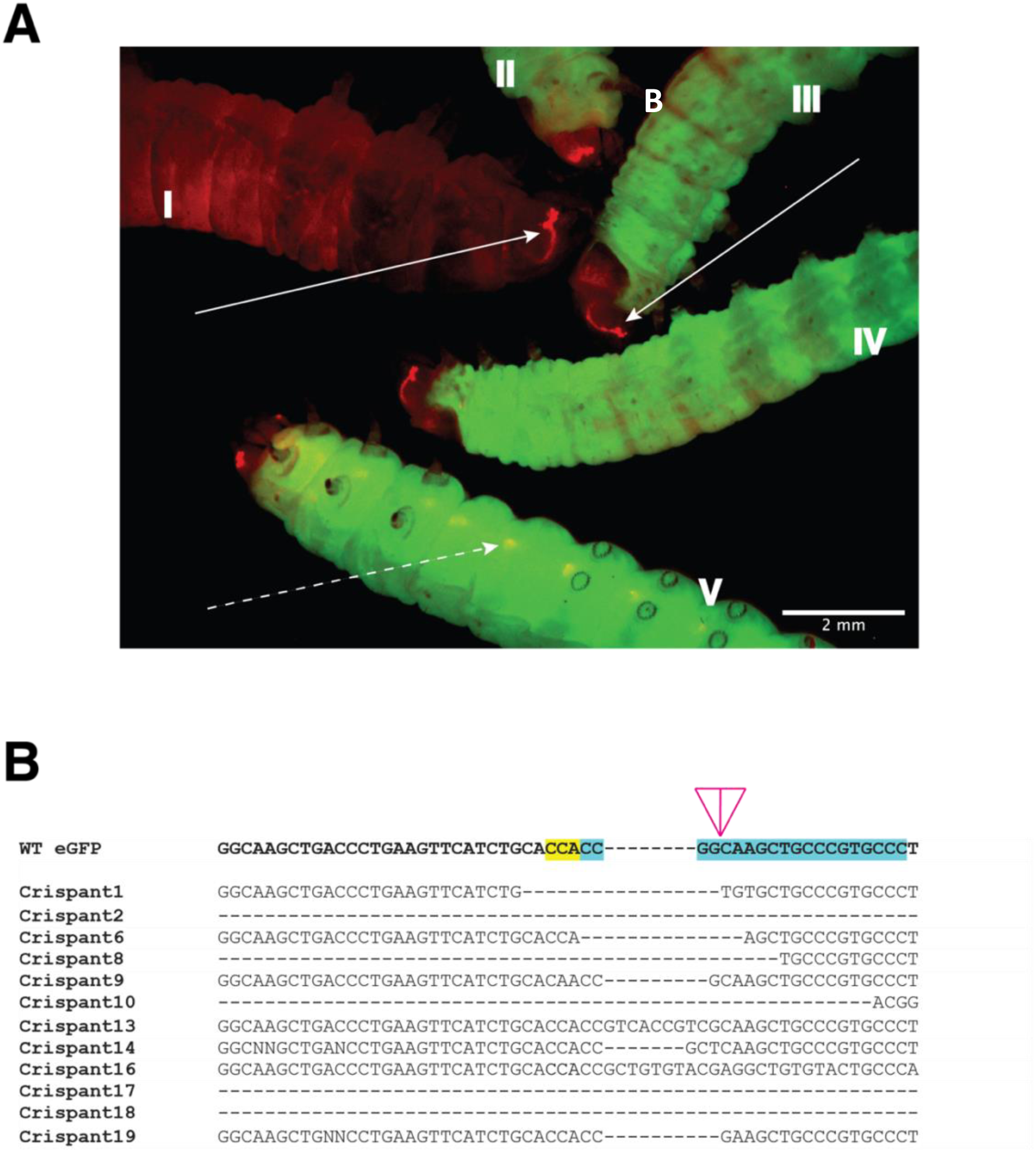
Demonastration of CRISPR in *Galleria* (A). *Bm*hsp90:GFP/3xP3:DsRed G0 larva I-V have an expression cassette inserted into their genome resulting in expression of eGFP visible in somatic un-melanised tissue and dsRed in both eye (solid arrow) and neural (dashed arrow) tissue. Larva I is the G1 offspring of individuals of the same transgenic line as II-V, but injected with a Cas9/sgRNA ribonucleoprotein targeting the eGFP gene, causing a heritable loss of function in the eGFP gene but retaining the eye specific dsRed expression. (B) Mutations within the coding sequences of 12 different GFP negative G1 crispants. The G1 larvae were offspring from individuals crosses of Bmhsp90:GFP/3xP3:DsRed G0s injected with an sgRNA targeting the N-terminus of the eGFP sequence, and crossed to wild type mates. The top line is the native eGFP sequence with the sequence complementary to the sgRNA highlighted in green and the PAM sequence highlighted in red.

## Discussion

The development of advanced genetic tools for model organisms such as rodents, *Drosophila* and zebrafish has revolutionised our ability to understand and interrogate their biology, shedding light on fundamental processes such as development, physiology and their response to environmental perturbations. Moreover, the capacity to genetically engineer these organisms to mimic human disease phenotypes has provided invaluable insights into the modes of action and potential efficacy of new therapeutic interventions. In this context, the establishment of methods for genetic modification of *Galleria mellonella* represents a transformative advancement with potentially profound implications for biomedical research.

Here, we have demonstrated the efficacy of both PiggyBac-mediated transformation for the tagging and expression of genes and gene knockout using CRISPR/Cas9 technology. Both techniques rely on the injection of exogenous nucleic acid into *Galleria* embryos, during the initial syncytial stage of development. Analysis of G0 pupae from multiple experiments, with different PiggyBac constructs, demonstrates the proportion of embryos hatching continuing through to pupation and eclosion varies between individual experiments and constructs and highlights areas for potential improvement. For instance, we have shown that different transposases can affect the transposition frequency; it is likely further optimisation of our methodology will result in increased rates. In addition, the spherical nature of *Galleria* embryos and the absence of phenotypic markers for the anterior-posterior axis precludes targeted microinjection to the germline precursor/pole cell region; an approach which has proved valuable in enhancing transgenesis efficiency in insects such as *Bombyx*^46^. Generating a transgenic *Galleria* line where the anterior/posterior pole of fertilised oocytes is highlighted (for example through fluorescent determinants, such as Nanos) could overcome this limitation although a Nanos promoter/terminator construct described in Heryanto et al ^47^ did not localise mScarlet to the presumptive germ region ^48^. Nonetheless, the PiggyBac donor integration rate using the methodology described here is generally at, or above, ∼1 % (Table 1), making the technique as it currently stands useful to researchers wishing to take up *Galleria* as a model.

The application of CRISPR/Cas9-mediated mutagenesis, now established in various insect species, represents another significant advancement. CRISPR/Cas9 offers a robust alternative to RNA interference (RNAi), which has been shown to work in *Galleria* ^49,50^, but with variable efficiency in other lepidoterans possibly due to compensatory gene upregulation ^51–53^. Again, although our current efficiencies are low, with further optimization of ribonucleoprotein concentrations and timing, survival rates and mutagenesis efficiencies in *Galleria* could potentially reach levels comparable to those reported in *Drosophila*, *Tribolium*, *Bombyx*, and other Lepidoptera.

Given that *Galleria* is predominantly used as a model to understand microbial infection and host-pathogen interactions, the methods described in this study have the potential to significantly enhance its utility and adoption. While this work focuses on the generation of reporter lines to visualise subcellular structures (the microtubule cytoskeleton and nuclei) future applications could include reporters of larval health status – for example, those driving fluorescence throughout the larvae, or in particular cell sub-types, upon infection by particular pathogens; or upon systemic release of anti-microbial peptides (AMPs). Such “sensor” moth larvae would provide a quantifiable read-out of health, complementary to current observation of melanisation. The ability to use CRISPR/Cas9 to knockout individual genes or entire gene families, or replace them with humanised disease variants, could also provide a bank of crispants that more accurately represent phenotypic traits found in human diseases, allowing screening for new therapeutics or interventions. Moreover, the ability to engineer *Galleria* not only broadens the scope of research and versatility of this model, but also aligns with the principles of the 3Rs (Replacement, Reduction, and Refinement). By offering a viable alternative or complementary system to rodent models, uptake of *Galleria* could lead to a reduction in the use of mammalian models in infection research and beyond, thus positively addressing both ethical and cost considerations of robust scientific animal research.

## Acknowledgments

We would like to Ivan Canada Luna for his help in maintenance of the GMRC *Galleria* laboratory colony, and for stimulating discussions. This work was funded by a PhD studentship jointly funded between the University of Exeter and Dstl (JCP), an NC3Rs Project Grant awarded to JGW (NC/T001518/1), which supported JSC, and an NC3R Training Fellowship, awarded to JCP (NC/W002388/1).

## Contributor Roles

JCP - conceptualisation, methodology, investigation, formal analysis, project administration, writing and funding acquisition (review & editing); JSC – methodology, investigation and writing (editing); JP – supervision, writing (editing), funding acquisition; RT: conceptualisation, resources, supervision, writing (editing); JGW - conceptualisation, methodology, project administration, supervision, writing (review & editing), funding acquisition.

## Disclosure of interest

The authors report no conflict of interest.

## Data availability statement

Raw data were generated at the University of Exeter Sequencing service and the University of Exeter Bioimaging Facility. Derived data supporting the findings of this study are available from the corresponding authors on request.

## Notes

### Competing Interest Statement

The authors have declared no competing interest.

